# Prenatal treatment with rapamycin restores enhanced hippocampal mGluR-LTD and mushroom spine size in a Down’s syndrome mouse model

**DOI:** 10.1101/2020.04.08.032029

**Authors:** Jesús David Urbano-Gámez, Itziar Benito, Juan José Casañas, María Luz Montesinos

**Affiliations:** Departamento de Fisiología Médica y Biofísica, Universidad de Sevilla, E-41009, Sevilla, Spain; Instituto de Biomedicina de Sevilla, IBIS/Hospital Universitario Virgen del Rocío/CSIC/Universidad de Sevilla. Sevilla, SPAIN; Servicio de Animalario, Hospital Universitario Virgen Macarena (HUVM), E-41009, Sevilla, Spain

**Keywords:** mGluR-LTD, mTOR, microelectrode array, MEA, dendritic spines, iTRAQ, proteomics, object recognition, trisomy 21, Down syndrome, Ts1Cje mice

## Abstract

Down syndrome (DS) is the most frequent genetic cause of intellectual disability including hippocampal-dependent memory deficits. We have previously reported hippocampal mTOR (mammalian target of rapamycin) hyperactivation, and related plasticity as well as memory deficits in Ts1Cje mice, a DS experimental model. Here we report that performance of Ts1Cje mice in novel object recognition (NOR) is impaired, but it is ameliorated by rapamycin treatment. Proteome characterization of hippocampal synaptoneurosomes (SNs) from these mice predicted an alteration of synaptic plasticity pathways, including long term depression (LTD), which was reversed by rapamycin. Accordingly, mGluR-LTD (metabotropic Glutamate Receptor-Long Term Depression) is enhanced in the hippocampus of Ts1Cje mice and this is correlated with an increased proportion of a particular category of mushroom spines in hippocampal pyramidal neurons. Remarkably, prenatal treatment of these mice with rapamycin normalized both phenotypes, supporting the therapeutic potential of rapamycin/rapalogs for DS intellectual disability.

## INTRODUCTION

DS, also known as trisomy 21, is one of the most common causes of intellectual disability. Among other difficulties, DS patients experience learning and memory deficits that evidence hippocampal dysfunctions (Pennington et al., 2003). We have previously shown that synaptic local translation, a key process for plasticity and hippocampal-dependent memory, is deregulated in the DS mouse model Ts1Cje (Sago et al., 1998) due to mTOR hyperactivation (Troca-Marín et al., 2011). Accordingly, mTOR hyperactivation has been also found in subjects with DS (Iyer et al., 2014; Perluigi et al., 2014). In Ts1Cje mice, mTOR hyperactivation seems to be caused by increased levels of Brain Derived Neurotrophic Factor (BDNF) that saturate the BDNF-TrkB-Akt-mTOR signaling axis (Troca-Marín et al., 2011; 2014). Furthermore, we also found that the BDNF-dependent Long Term Potentiation (LTP) is abolished in the Ts1Cje hippocampus, and that the specific mTOR inhibitor rapamycin fully restored this type of plasticity (Andrade-Talavera et al., 2015). Moreover, we observed that the impaired persistence of long-term memory (LTM) in the Barnes maze of Ts1Cje animals was also restored by rapamycin treatment (Andrade-Talavera et al., 2015).

The mTOR pathway is known to participate in other forms of plasticity, such as mGluR-LTD. This is mediated by group I metabotropic glutamate receptors and relies on protein translation (Huber et al., 2000). mTOR forms two different complexes: mTORC1, which contains, among other components, the defining protein RAPTOR (Regulatory Associated Protein of mTOR), and mTORC2, which contains RICTOR (Rapamycin-Insensitive Companion of TOR). In neurons, mTORC1 is mainly involved in translational control, mitochondrial function and autophagy regulation, whereas mTORC2 regulates actin cytoskeleton (reviewed by Costa-Mattioli and Monteggia, 2013). It is well known that rapamycin blocks hippocampal mGluR-LTD (Hou and Klann, 2004), initially suggesting that protein synthesis mediated by mTORC1 is necessary for this plasticity. Nevertheless, chronic treatment or high concentrations of rapamycin also inhibit mTORC2 (Sarbassov et al., 2006), and, moreover, it has been recently reported that mTORC2, but not mTORC1, is required for mGluR-LTD (Zhu et al., 2018). In line with these results, it is well established that inhibiting actin polymerization/depolymerization blocks mGluR-LTD (Zhou et al., 2011) and that dendritic spines elongate in response to mGluR activation, correlating with AMPA receptor (AMPAR) endocytosis (Vanderklish and Edelman, 2002; Zhou et al., 2011). In fact, mGluR activation induces rapid local translation of proteins involved in AMPAR internalization such as Arc/Arg3.1 (Activity Regulated Cytoskeleton-associated protein), OPHN1 (oligophrenin-1), MAP1B (Microtubule Associated Protein 1B) and STEP (Striatal-Enriched Protein Phosphatase) (Lüscher and Huber, 2010). These proteins are targets of FMRP (Fragile X Mental Retardation Protein, encoded by the *FMR1* gene), a key regulator of local translation. Since mGluR-LTD is enhanced in *FMR1* knockout mice, it has been proposed that FMRP serves as a brake on mGluR-stimulated protein synthesis (reviewed by Bhakar et al., 2012). Moreover, it has been shown that hippocampal mGluR-LTD requires the rapid synthesis and degradation of FMRP (Hou et al., 2006). Despite the extensive work, the definitive comprehension of the signaling pathways that contribute to protein synthesis necessary for mGluR-LTD remains elusive although roles for mTORC1 and ERK (Extracellular signal-regulated kinase) have been proposed (reviewed by Bhakar et al. 2012).

It has been suggested that hippocampal LTD has a specific role in encoding novelty and mGluR-LTD seems to be particularly important for spatial recognition of objects (Goh and Manahan-Vaughan, 2013a, 2013b). To gain insight into the possible use of rapamycin/rapalogs to improve cognition in the context of trisomy 21, we have analyzed the performance of untreated and rapamycin-treated Ts1Cje mice in the NOR test. We here report a clear impairment in performance of trisomic animals, compared to wild-type (WT) littermates, which was ameliorated by rapamycin treatment. By iTRAQ (isobaric tag for relative and absolute quantitation) we have characterized the proteome of hippocampal SNs from untreated and rapamycin-treated WT and Ts1Cje mice. Interestingly, mitochondrial function, calcium signaling and, remarkably, synaptic plasticity pathways, including LTD, were predicted to be altered in trisomic mice, but not in the rapamycin-treated siblings. Accordingly, we have found that the hippocampal mGluR-LTD is enhanced in Ts1Cje animals. Additionally, we have evaluated the effect of rapamycin on dendritic spine density and morphology. We found that prenatal treatment with rapamycin did not recover the decreased density of dendritic spines in Ts1Cje offspring but, interestingly, recovered the alterations observed in mushroom spine size. Strikingly, prenatal rapamycin treatment also normalized the mGluR-LTD in Ts1Cje hippocampus. Together, these results extend the evidence that supports the possible benefits of rapamycin for synaptic plasticity in the context of DS.

## MATERIALS AND METHODS

### Animals

Partial trisomic Ts1Cje mice (Sago et al., 1998) were purchased from Jackson Laboratories and, as recommended by the supplier, were maintained on a mixed genetic background by backcrossing Ts1Cje males to B6C3F1 hybrid females (supplied by Charles River). Sets of WT males and Ts1Cje littermates were used in the experiments. Blind animals homozygous for the retinal degeneration mutation *Pde6b*^rd1^, which segregates in the Ts1Cje colony (Sago et al., 1998), were not used. Experiments were carried out according to the European Union directive for the use of laboratory animals. All the protocols were approved by the Regional Government (Junta de Andalucía, Spain) Ethical Committee.

### Rapamycin treatment

For the treatment of adult mice, rapamycin (Seleckchem) was dissolved at 1 mg/ml in saline buffer containing 4% ethanol, 5% Tween®80 and 5% polyethylene glycol 400 (Kwon et al., 2003). A dose of 10 mg/kg was administered by intraperitoneal injection (one injection per day) during the 5 days prior to performance of object recognition tests or SNs preparation. For prenatal treatment, a single intraperitoneal injection of rapamycin (1 mg/kg) was applied to pregnant dams between E15 and E17 as previously described (Tsai et al., 2013). In this case rapamycin was prepared at 0,1 mg/ml in saline buffer with 1% ethanol, 0,25% Tween® 80, and 0.25% polyethylene glycol 400.

### Object recognition

Object recognition was performed as previously described (Leger et al., 2013). Briefly, 2-4 month-old male mice were subjected to a familiarization session in which two identical objects were placed 5 cm away from the walls inside a transparent box (40 cm × 40 cm surface, 35 cm height). Mice were free to explore both the right and the left objects for a total period of 20 s (or a maximum time of 10 min). As expected, no exploration preference was observed in this phase (not shown). The process was repeated 24 h later, but one of the familiar objects was replaced by a novel object. Mice were then allowed to freely explore the objects until they reached the criterion of exploring for a period of 20 s or otherwise, when the maximum session time (10 min) was reached. Animals that did not reach the criterion were excluded, as recommended (Leger et al., 2013). An automated tracking system (Smart, Harvard Apparatus) was used to register the behaviour of mice. To avoid olfactory cues, the box and the objects were cleaned with a surface disinfectant (1% CR36) after each trial. The experimenter was blind to the genotype of the animals. Memory performance was evaluated by comparing the mean time that mice spent exploring the new object with the chance level through a univariate t test, as indicated by Leger and cols. (2013).

### Synaptoneurosomes preparation

SNs were prepared as previously described (Troca-Marín et al., 2010). Briefly, hippocampi from 3 adult mice (2-4 month old) were rapidly dissected and homogenized in 12 ml of homogenization solution (10 mM Hepes, pH 7.4; 320 mM sucrose; 1.0 mM MgSO_4_; protease inhibitors leupeptin (10 μM), aprotinin (2 μg/ml), and pepstatin (1 μM)) with a glass-teflon Dounce homogenizer. The homogenate was centrifuged (1,000 g for 10 min at 4°C), and the pellet was subjected to an Optiprep discontinuous gradient (9-40% Optiprep). After centrifugation (16,500 g for 30 min at 4°C), the material from the first band (fraction O1) was recovered and subjected to discontinuous Percoll gradient (10-25%) centrifugation (32,400 g for 20 min at 4°C). The material from the fourth band (fraction 1P4), which is highly enriched in SNs, was recovered, washed with buffer, rapidly frozen and stored at −80°C until iTRAQ proteomics.

### iTRAQ labeling and analysis

Protein extraction, iTRAQ labeling and tandem mass spectrometry analysis was carried out at the Instituto de Biomedicina de Sevilla (IBiS) Proteomic Service. Briefly, synaptoneurosomal proteins were extracted using a lysis buffer that contained SDS, supplemented with protease inhibitors (Sigma), phosphatase inhibitor cocktails I and II (Sigma), and benzonase (Sigma). After incubation for 1 h, the samples were centrifuged for 15 min at 14,000 rpm in a refrigerated bench-top microfuge. Proteins present in the supernatant were quantified following iTRAQ labeling (AB ScieX) essentially following the manufacturer’s instructions, omitting the protein precipitation step in order to conserve minority proteins. 50 μg of proteins were labeled for each experimental condition in duplicates (8-plex). Data were analyzed using the Proteome Discoverer 1.4 software (Thermo), setting the False Discovery Rate (FDR) of both proteins and peptides identification to be less than 0.01.

### Pathway Analyses

Ingenuity Pathway Analysis (IPA, Fall release September 2019) was performed for both up- and downregulated proteins, considering a cutoff of 1.2-fold. This cutoff was established by analyzing the protein level variations observed in the WT sample replicate and determining the boundaries within the 80% best fitting replicate data.

The Ingenuity Knowledge Base (genes only) set was used as reference. All the molecules and/or relationships included in the analysis were experimentally observed, either in the mouse, rat, human nervous system tissue or neural (astrocytes or neurons) cells. The IPA software generates a list of significantly affected canonical pathways based on two parameters, the p-value and the Z-score. The p-value, calculated with the right-tailed Fisher Exact Test, is a measure of the probability that a particular pathway were enriched in the set of deregulated proteins due to random chance. To enhance the stringency of the analysis, we considered only pathways with a p-value ≤ 0.005 (i.e., −log_10_ (p-value) ≥ 2.3). Considering the protein level changes observed in the set of deregulated proteins, the canonical pathways identified are predicted to be either activated or inhibited applying the IPA Z-score algorithm. A positive Z-score value indicates that a pathway is predicted to be activated, and a negative Z-score indicates its inhibition. Canonical pathways which are not eligible for activity analysis by the IPA are marked as N/A.

### Antibodies

We used anti-FMRP (Abcam; Ref. ab69815), anti-MAP2 (Merck Millipore; Ref. MAB378), and anti-LC3B (Cell Signaling; Ref. #2775) as primary antibodies, while horseradish peroxidase (HRP)-conjugated anti-rabbit (Promega), Alexa Fluor 555 goat anti-rabbit (Invitrogen), and Alexa Fluor 488 goat anti-mouse (Invitrogen) were used as secondary antibodies.

### Western blot

Western blot was performed as previously described (Casañas et al., 2019). Briefly, total hippocampal protein extracts were prepared by mechanical tissue homogenization in extraction buffer containing SDS and protease and phosphatase inhibitor cocktails. Protein extracts were resolved by SDS-PAGE on Mini-PROTEAN® TGX Stain-Free™ precast gels (BioRad). Proteins were transferred to polyvinylidene difluoride (PVDF) membranes. HRP-conjugated secondary antibodies were detected with the WesternBright Quantum HRP Substrate (Advansta) and chemiluminiscence measured on a ChemiDoc XRS (BioRad) imager.

### Immunocytochemistry

Hippocampal cultures from postnatal day (P) 0 WT or Ts1Cje littermates were established as previously described (Alves-Sampaio et al., 2010). After 14 days in vitro (DIV), cells were fixed with 4% paraformaldehyde (PFA) in phosphate-buffered saline (PBS) and subjected to dual immunocytochemistry using antibodies against FMRP and MAP2. Confocal microscopy images were acquired and processed as previously described (Casañas et al., 2019). The mean pixel intensity for the corresponding immunofluorescent FMRP signal in dendrites was determined (in arbitrary units, a.u.) using a Matlab (Mathworks) routine previously established (Troca-Marín et al., 2011).

### Electrophysiological recordings

Hippocampal slices were prepared from P21-P30 WT and Ts1Cje male mice. Mice were anesthetized and brains were removed in an ice-cold artificial cerebrospinal fluid with partial substitution of Na with sucrose (ACSFs) with constant flux of carbogen (5% CO_2_; rest O_2_; H_2_O <5 ppm). ACSFs composition was: 2.5 mM KCl; 1.25 mM NaH_2_PO_4_ (2H_2_O); 26 mM NaHCO_3_; 25 mM glucose; 0.5 mM CaCl_2_ (2H_2_O); 4 mM MgSO_4_ (7H_2_O); 185 mM sucrose). Brains were positioned in the cutting chamber over a thin film of ethyl cyanoacrylate and were submerged in cold ACSFs. 350 μm horizontal slices were obtained using a vibratome (Vibratome 3000, Sectioning System) and both hippocampi were isolated in each slice preserving entorhinal cortex adjacent to the hippocampal formation. Slices were incubated in ACSF (126 mM NaCl; 3 mM KCl; 1.3 mM MgSO_4_ (7H_2_O); 2 mM CaCl_2_ (2H_2_O); 1.25 mM NaH_2_PO_4_ (2H_2_O); 24 mM NaHCO_3_; 10 mM glucose) for 30 minutes at 34°C. Then, slices were incubated for at least 2 hours in ACSF at room temperature (RT; 22-25°C) before recording, keeping constant the flux of carbogen. Slices were transferred to a 3D-MEA chamber (MultiChannel Systems, Ref.60-3DMEA200/12/50iR-gr) and stayed at least 10 min before stimulation with a 2 ml/min flux rate of ACSF. All the registers were performed at RT. The experimenter was blind to the genotype of the animals.

The 3D-MEA devices content 59 conical TiN electrodes with 12 μm diameter (100 μm in the base of the electrode) and 50 μm high, distributed in an 8×8 matrix with an internal reference electrode at the position 15 (column 1, row 5). The inter-electrode distance is 200 μm. The isolation material used for these devices’ circuitry is SiN and the base is made out of glass.

Slices were stimulated at the Schaffer collateral pathway in the CA1 region of hippocampus using a biphasic square pulse (negative phase-positive phase; 100 ms per phase) at 0.0167 Hz (1 stimulus per minute). Slices showing field excitatory postsynaptic potential (fEPSP) amplitude lower than 100 μV for test stimulation (−/+1,750 mV) in the CA1 region were discarded. Input/output (I/O) curve was performed reaching the limits of the system (750 mV-4,000 mV; 3 stimuli per amplitude) and baseline was stablished at 60% of the highest response obtained at the maximum voltage applied. After baseline stabilization, 100 μM (S)-3,5-Dihydroxyphenylglycine (DHPG; Sigma-Aldrich) in ACSF was bath applied at a flow rate of 2 ml/min for 5 minutes. DHPG-long term depression (DHPG-LTD) was recorded for 1 hour after treatment. In order to determine the synaptic efficacy, the slope of the initial fEPSP curve was measured in the segment comprised between the 10-90% of the curve amplitude. Recordings were acquired at 10,000Hz.

Electrophysiological results are presented as the mean +/− SEM normalized to the baseline slope mean. For statistical analysis, a t-Student test was performed for compared experimental groups. Normalized slope at the minute 60 after DHPG treatment initiation was used for comparisons.

### Golgi staining and spine morphology analysis

WT and Ts1Cje P18 mice were anesthetized and both hemispheres were separated, and the cerebellum was removed. A FD Rapid Golgi stain kit (FD NeuroTechnologies INC) was used following the manufacturer guidelines. After the staining process, 100 μm coronal slices were prepared using a cryostat (Leica CM1950). Slices were deposited over Menzel-Gläser Superfrost PLUS microscope slides (ThermoScientific) and covered with a thin gelatine coat (Gelatine Gold; Panreac DC).

Secondary dendrites images were acquired from the hippocampal CA1 stratum radiatum region, at a 1,600 × 1,200 pixel resolution, using an Olympus BX61 microscope (CellSens software). For spine morphology analysis, image acquisition was made using an UPlanSApo60x/1.35 oil objective, with a 10x ocular and an additional 2x magnification. For spine density analysis, images were acquired using an UPlanApo40x/0.85 air objective, with a 10x ocular, and an additional 2x magnification. To determine dendritic spine density, spines were manually counted, and the branch longitude was measured using ImageJ software. For spine morphology analysis, spines perimeters were manually defined, and the head diameter was measured using ImageJ software.

The experimenter was blind to sample genotype and treatment.

## RESULTS

### Memory NOR test is impaired in Ts1Cje mice and is slightly improved by rapamycin treatment

In a previous work we found that Ts1Cje mice showed neither learning nor memory deficits in the Barnes maze test. Nevertheless, these mice presented impaired LTM persistence, which was restored by rapamycin (Andrade-Talavera et al., 2015). To further characterize the behavior of this DS mouse model, we performed a NOR test.

Four experimental groups were established: vehicle-treated WT or Ts1Cje mice, and rapamycin-treated WT (WT RAPA) or Ts1Cje (Ts1Cje RAPA) mice (see Materials and Methods). WT mice showed a memory index = 72.6%, significantly higher than the chance level (i.e., 50%) indicating that NOR memory was normal in WT animals (Fig. 1). On the other hand, the memory of Ts1Cje mice in NOR was clearly impaired (memory index = 43.8%). Remarkably, rapamycin had a negative effect on NOR performance in WT animals (memory index = 44.5%), whereas it seemed to have a slightly positive effect in Ts1Cje mice (memory index = 57.7%), although this value was not significantly different to the chance level. Nevertheless, a two-tailed Mann-Whitney test comparison between the memory indexes of Ts1Cje and Ts1Cje RAPA groups showed a p-value = 0.09, which is near to significance, suggesting a modest positive effect of rapamycin on Ts1Cje NOR memory.

**Figure 1.**
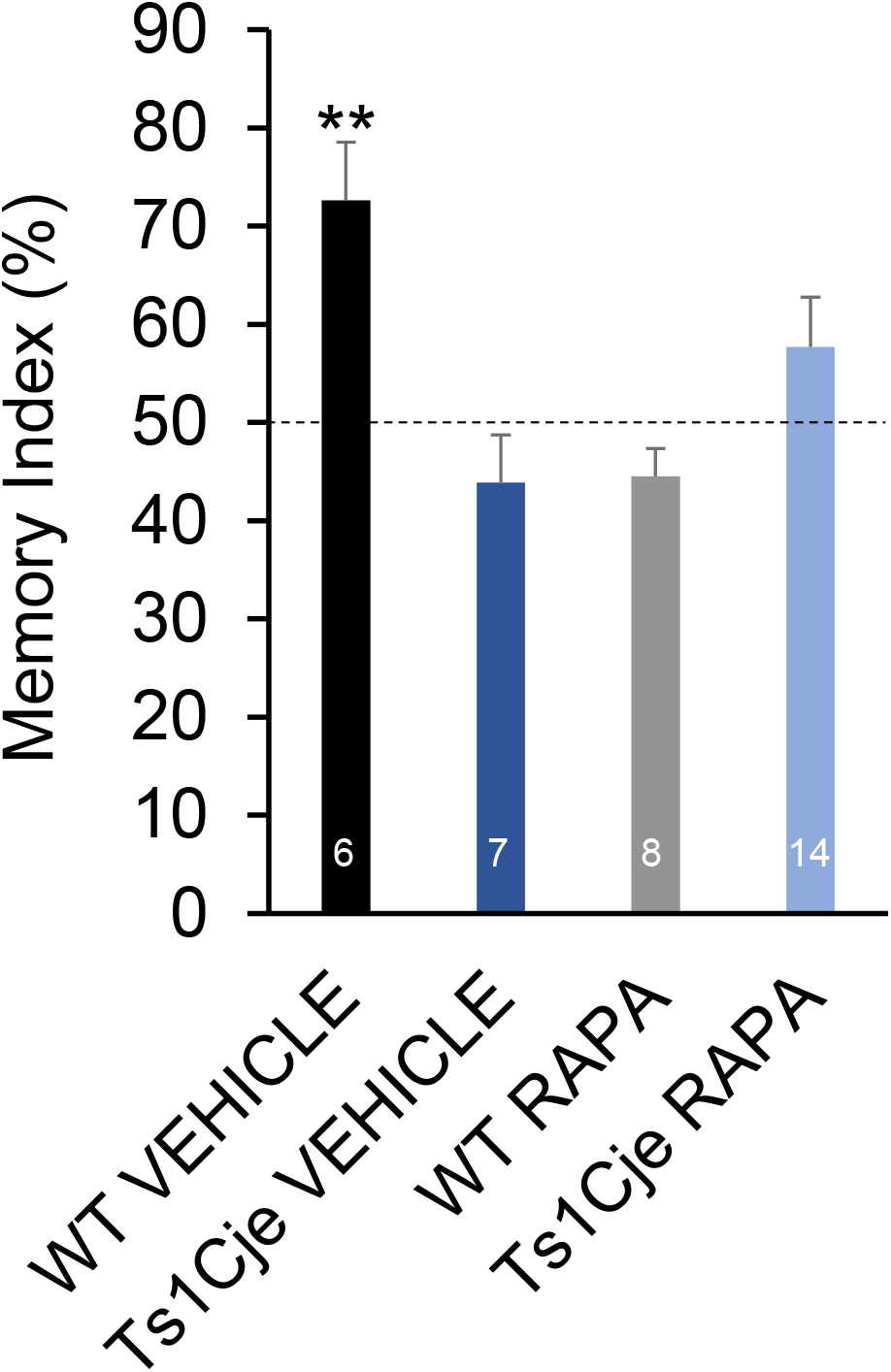
Novel object recognition (NOR) test in vehicle and rapamycin-treated WT and Ts1Cje mice. Animals injected with vehicle or rapamycin (RAPA) were subjected to the NOR test. Memory index is shown as the percentage of time mice spent exploring the new object (total exploration time 20 s). The dashed line indicates the chance level of performance (50%). One-sample t-test for the mean different from 50% showed a significant p value only in the case of WT vehicle as indicated with asterisks (p = 0.012, n = 6 for WT vehicle; p = 0.26, n = 7 for Ts1Cje vehicle; p = 0.09, n = 8 for WT RAPA; p = 0.145, n = 14 for Ts1Cje RAPA). Data are expressed as the mean ±SEM. The number of animals for each condition is indicated in the corresponding bar.

### Proteomic analysis of hippocampal SNs from Ts1Cje and rapamycin-treated mice

In order to identify synaptic differences that could account for plasticity and memory deficits in Ts1Cje mice and to evaluate the effect of rapamycin treatment on their synaptic proteome, we performed a proteomic analysis of hippocampal SNs isolated from WT and Ts1Cje mice, treated or not with rapamycin. As in the NOR test, rapamycin was administered by a daily intraperitoneal injection along the 5 days prior to SNs isolation.

Protein samples were subjected to 8-plex iTRAQ proteomics. Technical replicates of the four biological samples were included, and the relative amount of each detected protein was referenced to one of the WT samples (Table S1 in Supplementary File 1). 1,890 proteins were identified by at least two unique peptides (Table S2 in Supplementary File 1). Proteins not detected in all the samples (4 proteins) as well as technical replicates with CV% >30% (19 proteins) were removed. The geometric mean of the relative protein amount was calculated (Table S3 in Supplementary File 1) and used in the subsequent analysis by IPA software. From the 1,867 proteins initially included in the IPA, 12 resulted to be unmapped, and from the 1,855 mapped identities, IPA considered 1,824 of them as “analysis-ready”.

Considering a 1.2-fold cutoff (see Materials and Methods), 116 proteins were found to be deregulated (108 up- and 8 down-regulated) in Ts1Cje SNs compared to WT SNs (Table S4 in Supplementary File 2). In SNs from the Ts1Cje RAPA group 88 proteins were affected (58 up- and 30 down-regulated) (Table S5 in Supplementary File 2) while in the WT RAPA group 130 proteins were deregulated (104 up- and 26 down-regulated), compared to WT SNs (Table S6 in Supplementary File 2). A Venn diagram representation (Fig. 2) showed 58 proteins similarly affected in WT RAPA and Ts1Cje RAPA SNs, suggesting that these proteins were affected by rapamycin regardless the genotype.

**Figure 2.**
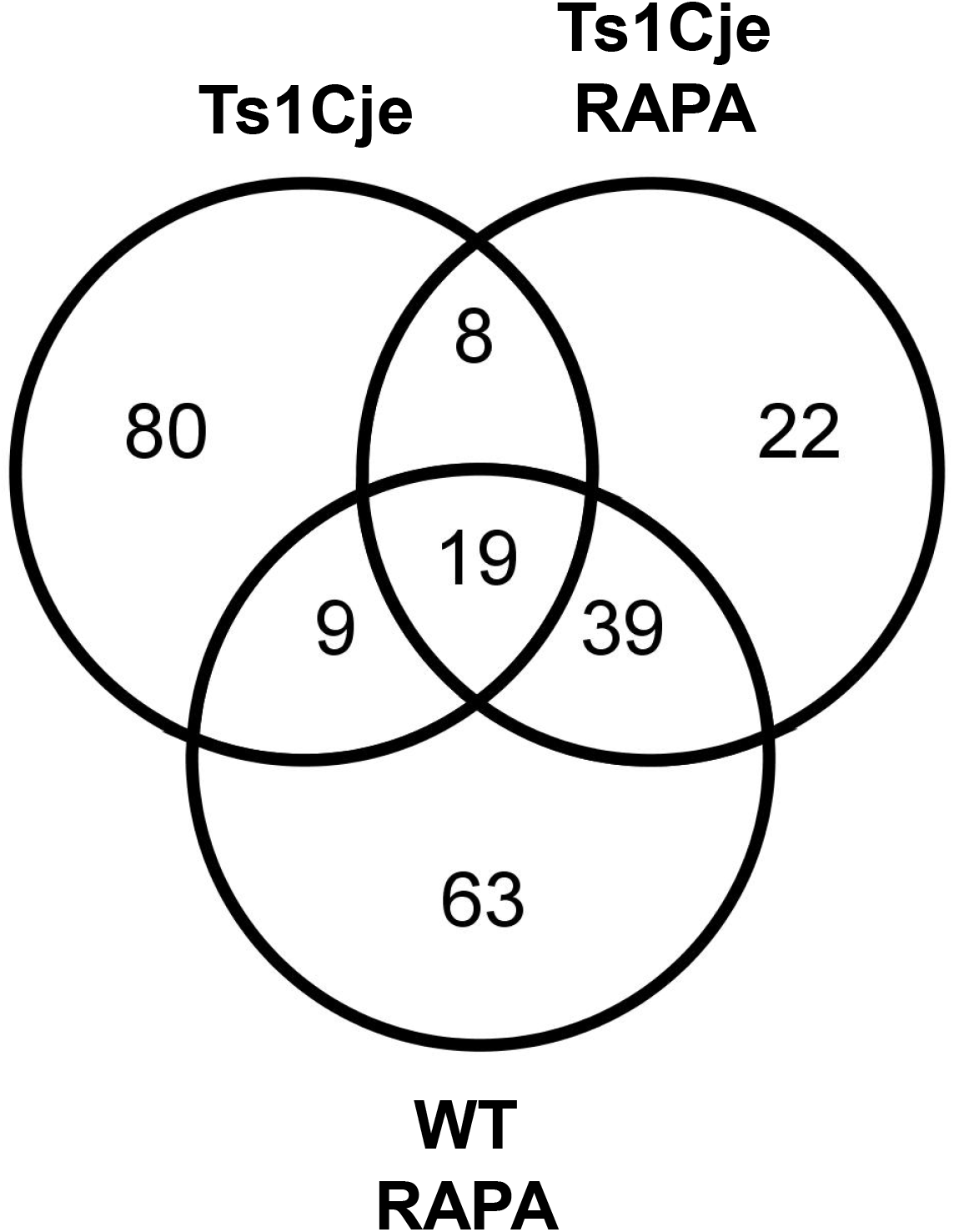
Venn diagram of proteins deregulated in hippocampal SNs from Ts1Cje mice and from rapamycin-treated Ts1Cje and WT mice. The numbers of overlapping proteins among the experimental groups are shown.

Attending to the Fischer p-value, IPA revealed that, the mitochondrial dysfunction canonical pathway, and the partially overlapping oxidative phosphorylation and sirtuin signaling pathways were among the most significantly enriched in the set of proteins affected in Ts1Cje SNs (Fig. 3 and Table S7 in Supplementary File 3).

**Figure 3.**
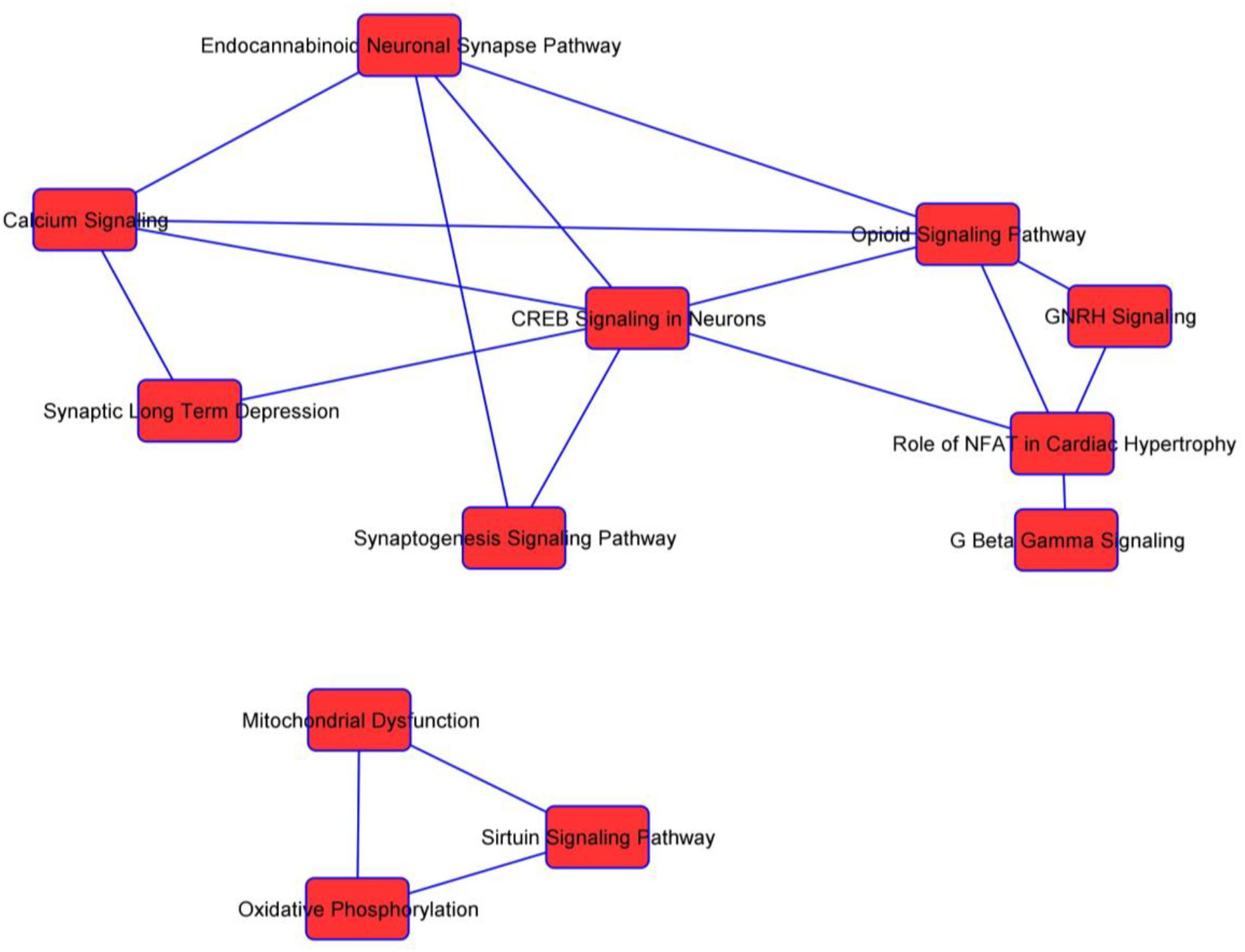
Canonical IPA pathways affected in Ts1Cje SNs. The most significant canonical pathways identified by IPA among the altered proteins in Ts1Cje SNs are shown. Overlapping pathways sharing at least 7 proteins are connected by solid lines.

Accordingly, mitochondrial alterations have been described in DS and DS mouse models (reviewed by Valenti et al., 2014; 2018; Izzo et al., 2018). In fact, it has been recently shown that reduced autophagy/mitophagy due to mTOR hyperactivation produces damaged mitochondria accumulation in DS fibroblasts (Bordi et al., 2019). In agreement with these data, we have found reduced levels of the B-II isoform of Microtubule-associated protein 1A/1B-light chain 3 (LC3B-II), a marker of autophagy, in Ts1Cje hippocampus, compared to WT (Fig. 4). Remarkably, when we compared the relative amounts of proteins involved in mitochondria-related pathways in the different experimental groups (Table 1) we found that the levels of these proteins were normal in Ts1Cje RAPA SNs, compared to WT, suggesting that the rapamycin treatment rescued the mitochondrial Ts1Cje phenotype.

**Table 1.**
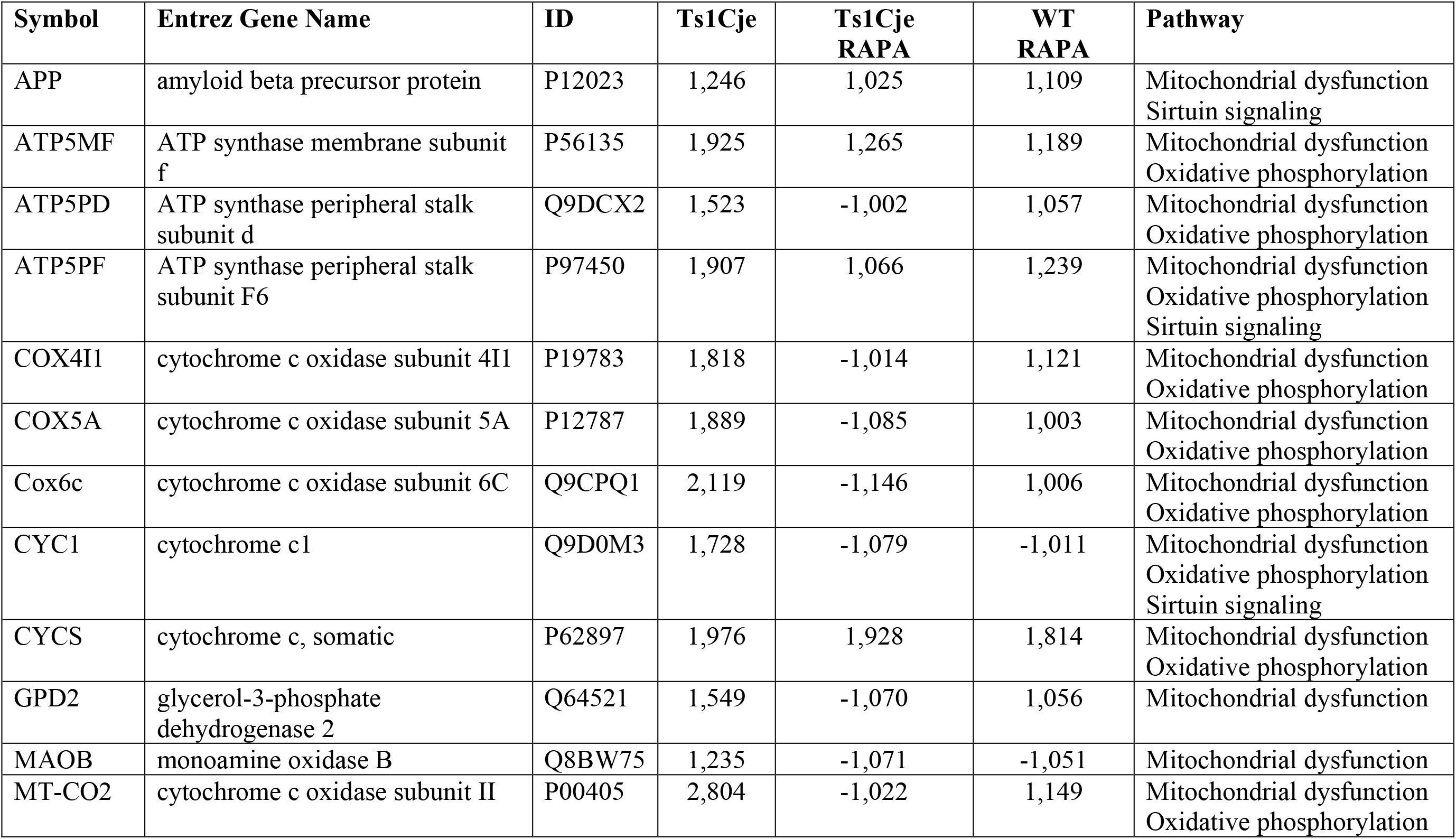
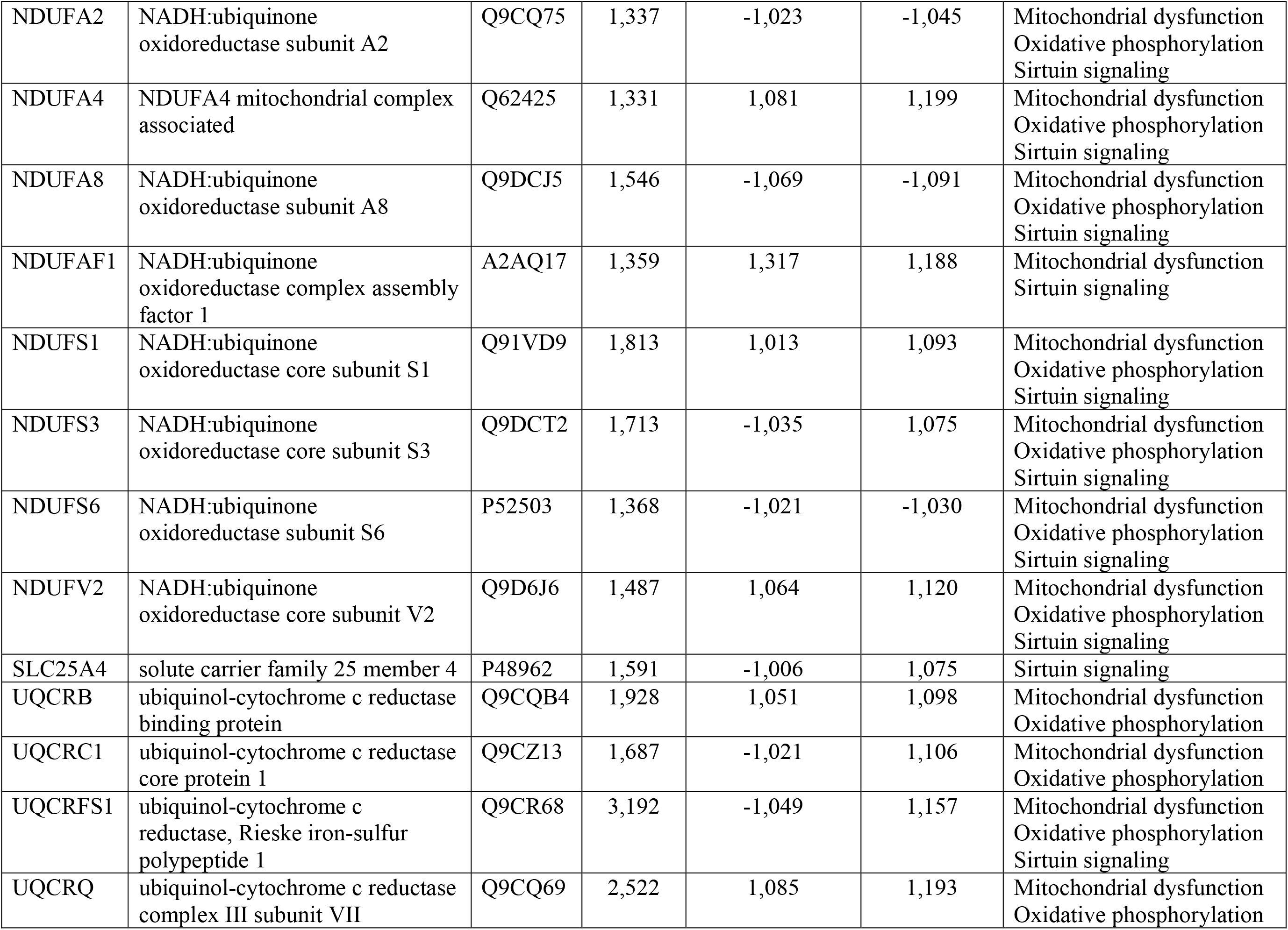
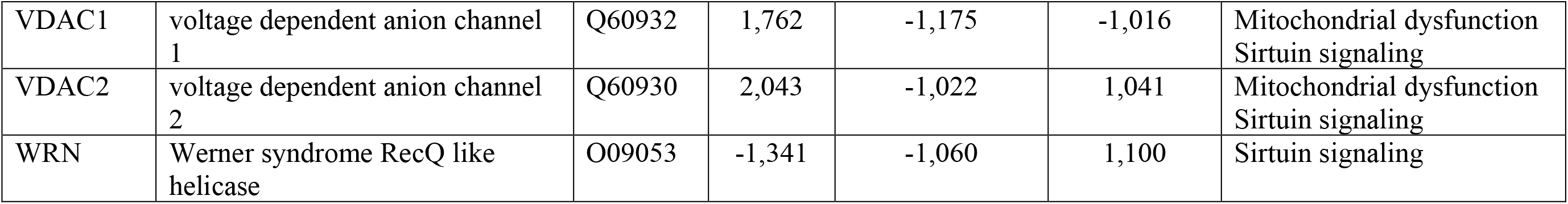
Proteins involved in mitochondrial pathways affected in Ts1Cje SNs. The relative amounts (fold change, compared to WT SNs) of these proteins in Ts1Cje, Ts1Cje RAPA and WT RAPA SNs are shown.

**Figure 4.**
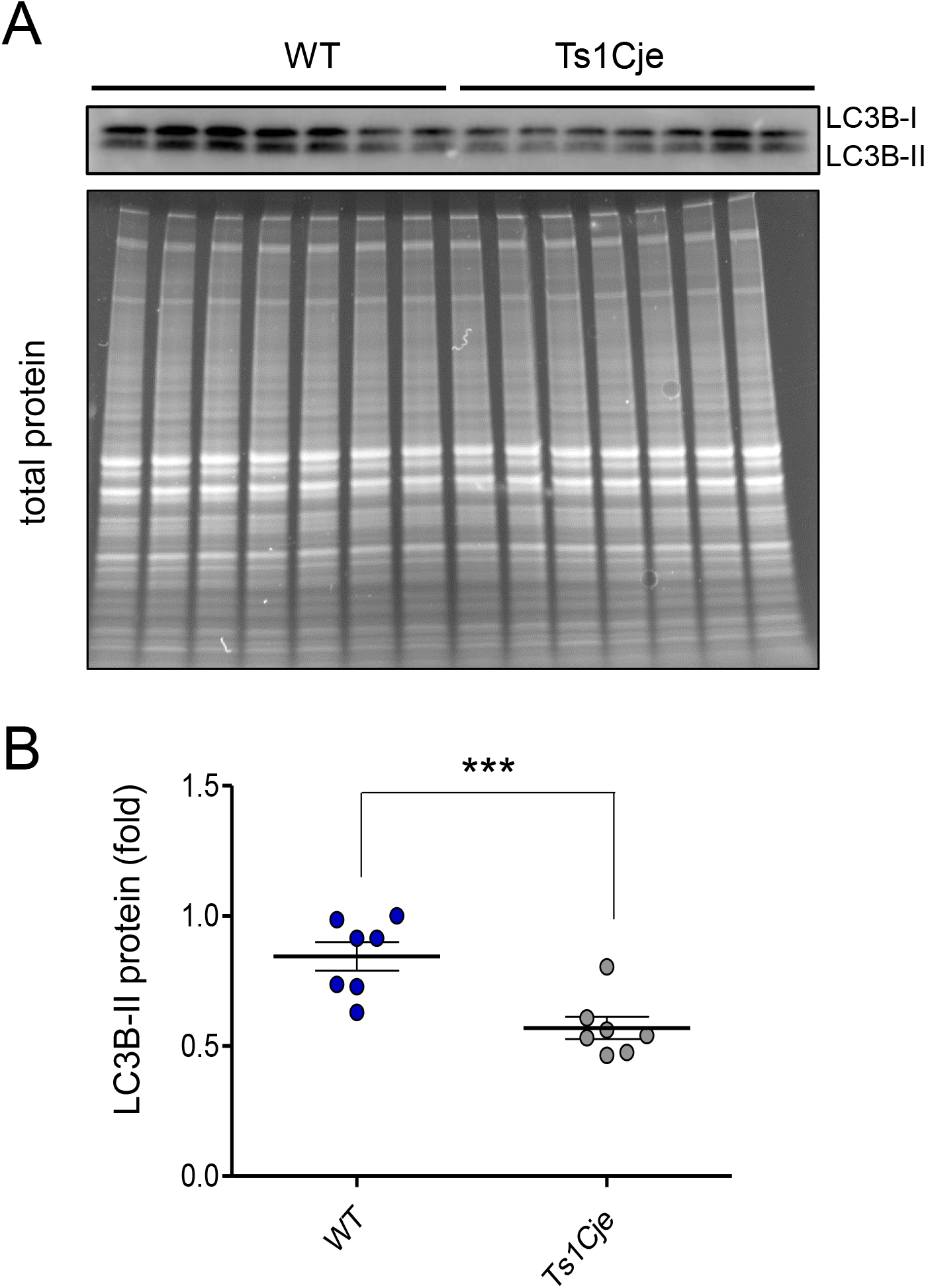
Quantification of LC3B-II protein in WT and Ts1Cje hippocampus. Hippocampal proteins from WT and Ts1Cje mice pairs were analyzed in Western blots with anti-LC3B antibody. (A) Western blot showing WT and Ts1Cje littermate pairs analyzed. (B) Quantification. The signals were normalized to the corresponding total protein loaded and the mean ± SEM values are shown (p = 0.0041, Mann Whitney test, n = 7).

Most important for the objectives of this work, we found that some key synaptic plasticity related pathways, including calcium signaling, CREB signaling, the endocannabinoid neuronal synapse pathway, the glutamate receptor signaling, and the synaptic LTD, were also significantly affected in Ts1Cje SNs (Fig. 3 and Table S7 in Supplementary File 3). These data are detailed in Table 2 for the different experimental groups, compared to WT. Very interestingly, the Z-score analysis predicted increased activity of these pathways in Ts1Cje SNs, but not in Ts1Cje RAPA (Table S8 in Supplementary File 3), suggesting that the rapamycin treatment also normalized the synaptic levels of proteins involved in these synaptic functions.

**Table 2.** Proteins involved in synaptic plasticity pathways affected in Ts1Cje SNs. The relative amounts (fold change, compared to WT SNs) of these proteins in Ts1Cje, Ts1Cje RAPA and WT RAPA SNs are shown.

| Symbol | Entrez Gene Name | ID | Ts1Cje | Ts1Cje RAPA | WT RAPA | Pathway |
| --- | --- | --- | --- | --- | --- | --- |
| ADCY9 | adenylate cyclase 9 | P51830 | 1,344 | 1,080 | 1,210 | CREB signaling<br>Endocannabinoid neuronal synapse |
| ATP2A2 | ATPase sarcoplasmic/endoplasmic reticulum Ca<sup>2+</sup> transporting 2 | O55143 | 1,251 | -1,033 | 1,004 | Calcium signaling |
| ATP2B1 | ATPase plasma membrane Ca<sup>2+</sup> transporting 1 | G5E829 | 1,329 | -1,077 | 1,029 | Calcium signaling |
| ATP2B2 | ATPase plasma membrane Ca<sup>2+</sup> transporting 2 | F8WHB1 | 1,268 | 1,017 | 1,043 | Calcium signaling |
| CACNA1A | calcium voltage-gated channel subunit alpha1 A | P97445 | 1,314 | 1,117 | -1,007 | Calcium signaling<br>CREB signaling<br>Endocannabinoid neuronal synapse<br>Synaptic LTD |
| CACNA1G | calcium voltage-gated channel subunit alpha1 G | Q5SUF8 | 1,474 | -1,164 | -1,029 | Calcium signaling<br>CREB signaling<br>Endocannabinoid neuronal synapse<br>Synaptic LTD |
| CACNA2D1 | calcium voltage-gated channel auxiliary subunit alpha2delta 1 | O08532 | 1,262 | -1,117 | -1,034 | Calcium signaling<br>CREB signaling<br>Endocannabinoid neuronal synapse<br>Synaptic LTD |
| GNB2 | G protein subunit beta 2 | E9QKR0 | 1,207 | -1,033 | 1,092 | CREB signaling |
| GNG2 | G protein subunit gamma 2 | P63213 | -1,212 | -1,319 | -1,404 | CREB signaling<br>Endocannabinoid neuronal synapse<br>Glutamate receptor signaling |
| GRIA2 | glutamate ionotropic receptor<br>AMPA type subunit 2 | P23819 | 1,481 | 1,132 | 1,173 | Calcium signaling<br>CREB signaling<br>Endocannabinoid neuronal synapse<br>Glutamate receptor signaling<br>Synaptic LTD |
| GRIA4 | glutamate ionotropic receptor<br>AMPA type subunit 4 | Q9Z2W8 | -1,251 | -1,142 | -1,170 | Calcium signaling<br>CREB signaling<br>Endocannabinoid neuronal synapse<br>Glutamate receptor signaling<br>Synaptic LTD |
| GRIN1 | glutamate ionotropic receptor<br>NMDA type subunit 1 | A2AI21 | 1,758 | 1,050 | 1,129 | Calcium signaling<br>CREB signaling<br>Endocannabinoid neuronal synapse<br>Glutamate receptor signaling |
| GRIN2B | glutamate ionotropic receptor<br>NMDA type subunit 2B | Q01097 | 1,371 | 1,031 | -1,030 | Calcium signaling<br>CREB signaling<br>Endocannabinoid neuronal synapse<br>Glutamate receptor signaling |
| GRM5 | glutamate metabotropic<br>receptor 5 | Q3UVX5 | 1,200 | 1,012 | -1,045 | CREB signaling<br>Endocannabinoid neuronal synapse<br>Glutamate receptor signaling<br>Synaptic LTD |
| RAP2B | RAP2B, member of RAS<br>oncogene family | P61226 | 1,300 | 1,029 | 1,144 | Calcium signaling<br>CREB signaling<br>Synaptic LTD |
| RYR2 | ryanodine receptor 2 | E9Q401 | 1,251 | 1,043 | -1,045 | Calcium signaling<br>Synaptic LTD |

### mGluR-LTD is enhanced in the CA1 region of Ts1Cje hippocampus

As mentioned above, the IPA analysis suggested that LTD is increased in the Ts1Cje hippocampus and, moreover, we found impaired NOR in Ts1Cje mice (Fig. 1).

Interestingly, novelty and spatial recognition of objects engage mGluR-LTD (Goh and Manahan-Vaughan, 2013a, 2013b) It is also well known that mTOR is involved in this type of plasticity (Hou and Klann, 2004; Zhu et al., 2018) and, remarkably, mTOR is hyperactivated in Ts1Cje hippocampus (Troca-Marín et al., 2011). Altogether, these data stimulated us to evaluate mGluR-LTD in Ts1Cje mice.

mGluR-LTD is experimentally induced in hippocampal slices by a brief (5 min) exposure to DHPG, a specific agonist of mGluR1/5 receptors (Huber et al., 2000). As expected, an LTD of evoked excitatory synaptic responses was evident in WT slices after DHPG exposure (Fig. 5, 82.12 ± 8.31% of baseline, n = 8 slices from 4 mice). Remarkably, an enhanced mGluR-LTD was elicited in Ts1Cje hippocampal slices (Fig. 5, 59.12 ± 4.92% of baseline, n = 7 slices from 3 mice). In conclusion, we found that mGluR-LTD was exaggerated in the Ts1Cje CA1 hippocampal region.

**Figure 5.**
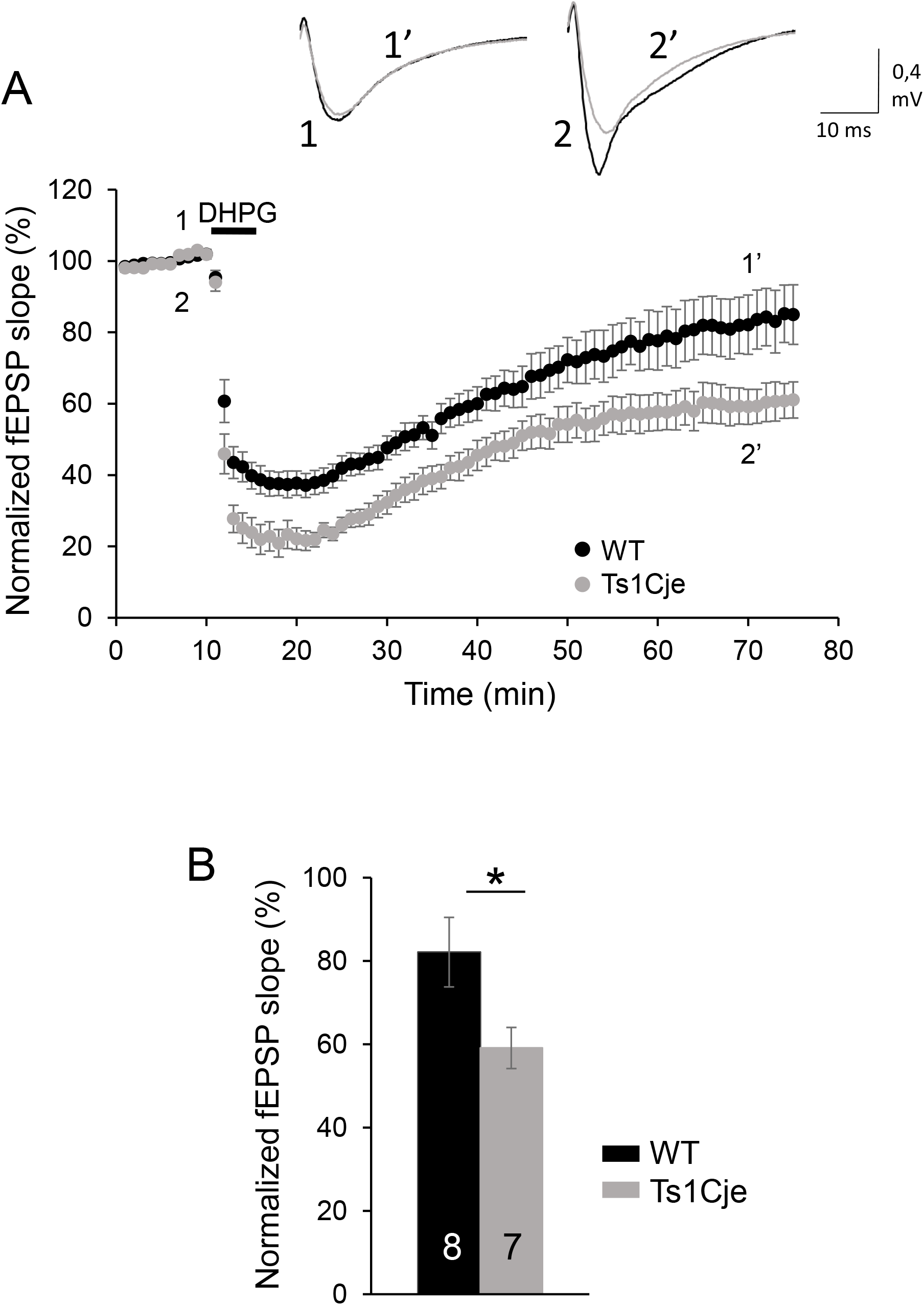
mGluR-LTD in WT and Ts1Cje hippocampal slices. (A) Time course of DHPG effects on field excitatory postsynaptic potentials (fEPSP) in WT and Ts1Cje mice. Upper insets: representative traces of a fEPSP before (1, 2) and after (1’, 2’) DHPG application in WT (1, 1’) and Ts1Cje (2, 2’) mice. (B) Quantification of the effects depicted in panel A. The error bars represent the SEM. The number of slices for each condition is indicated in the corresponding bar. P = 0.039 Student’s t-test.

Enhanced mGluR-LTD is a hallmark of *FMR1* knockout mice (Nosyreva and Huber, 2006). As already mentioned, FMRP (the *FMR1* encoded protein) is a repressor of local synthesis of proteins necessary for mGluR-LTD (Park et al., 2008; Nadif Kasri et al., 2011; Niere et al., 2012). Thus, we evaluated the amounts of FMRP in Ts1Cje hippocampus by Western blot and, strikingly, we found a slight, yet significant, increase of FMRP (Fig. 6A and B). To evaluate more precisely the amount FMRP in the dendritic compartment, we performed double immunocytochemistry in primary cultures of hippocampal neurons at DIV 14 and measured the fluorescence level of FMRP labeling in MAP2-positive neurites (i.e., dendrites). As shown in Fig. 6C, D, dendritic FMRP labeling was 1.6-fold higher in Ts1Cje neurons than in those from the WT. Hence, we must conclude that despite the higher levels of FMRP, mGluR-LTD was abnormally enhanced in Ts1Cje hippocampus.

**Figure 6.**
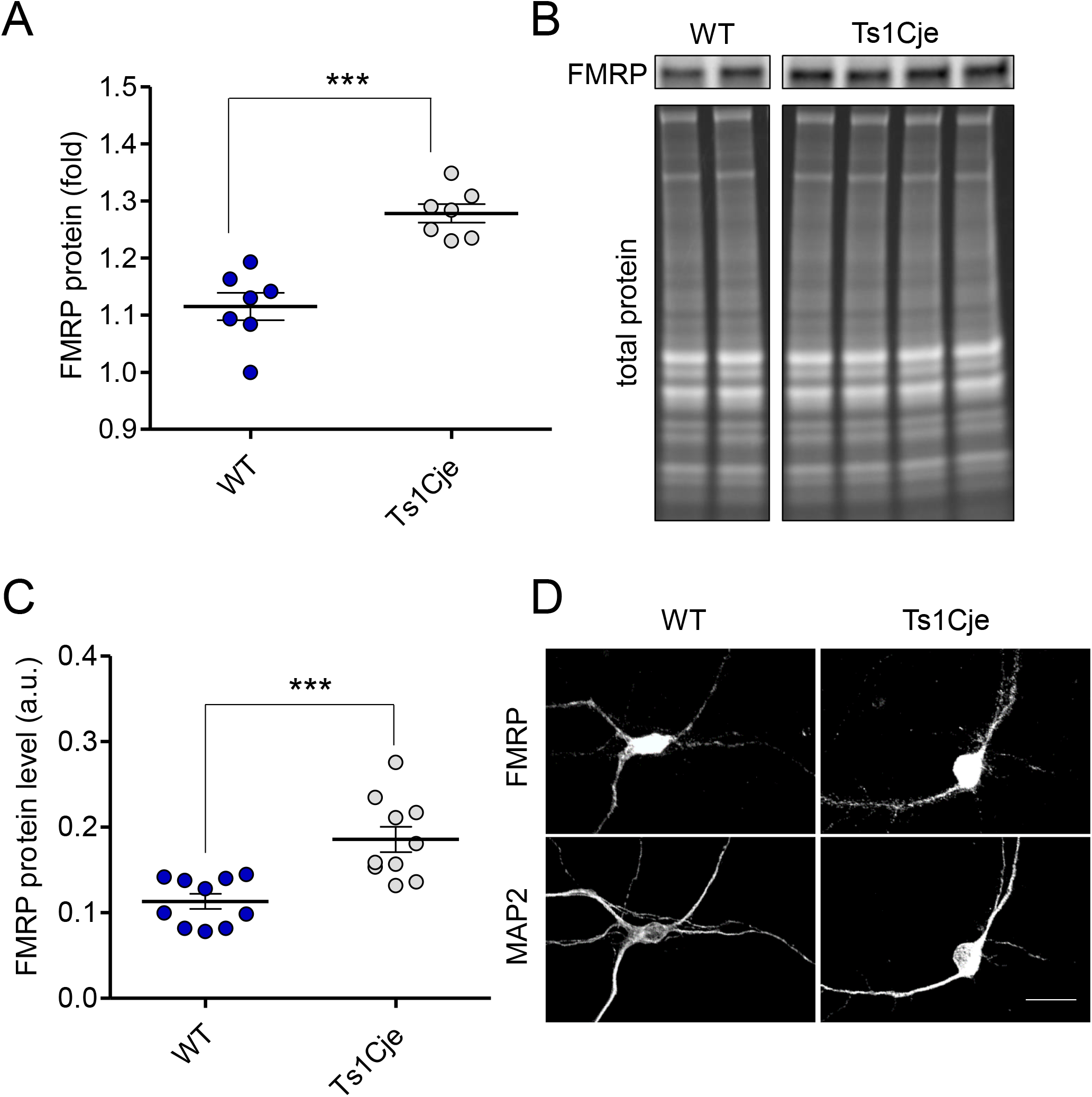
Quantification of FMRP protein in WT and Ts1Cje hippocampus. (A) Hippocampal proteins from WT and Ts1Cje mice pairs were analyzed by Western blots with an anti-FMRP antibody. The signals were normalized to the corresponding total protein loaded and the mean ± SEM values are shown (p = 0.0006, Mann Whitney test, n = 7). (B) Representative Western blot showing two WT and four Ts1Cje littermate pairs (some lanes corresponding to non-littermate animals have been removed). (C) Quantification of the relative amount of FMRP protein in dendrites of WT and Ts1Cje hippocampal neurons at DIV14. The mean pixel intensity ± SEM for FMRP immunofluorescence in dendrites is shown in arbitrary units (a.u.) (p = 0.0006, t test, n = 10). (D) Representative gray scale confocal images from the experiment in panel C, showing FMRP and MAP2 labeling in WT and Ts1Cje neurons. Scale bar = 20 μm. Note that dendritic branching and length are reduced in Ts1Cje neurons, compared to WT controls (unpublished observations).

### Prenatal treatment with rapamycin normalizes size distribution of mushroom type dendritic spines in the stratum radiatum of Ts1Cje mice

It has been proposed that LTD is a physiological correlate of synapse pruning (Nägerl et al., 2004; Zhou et al., 2004; Sheng and Erturk, 2014). Since dendritic spine alterations have been described in DS and DS mouse models including Ts1Cje (Benavides-Piccione et al., 2004; Belichenko et al., 2004, 2007), we decided to evaluate the effect of prenatal treatment with rapamycin on dendritic spine density in postnatal Ts1Cje mice using a rapid Golgi stain method (see Materials and Methods). We observed a reduced spine density in secondary dendrites of CA1 stratum radiatum of untreated Ts1Cje animals, compared to WT controls (WT: 0.640 ± 0.031 spines/μm, n = 50; Ts1Cje: 0.523 ± 0.035 spines/μm, n = 26; t-Student test p-value = 0.015). However, prenatal treatment with rapamycin produced no significant effect on spine density in either group (WT RAPA: 0.666 ± 0.029 spines/μm, n = 27; Ts1Cje RAPA: 0.572 ± 0.021 spines/μm, n = 31).

It has been shown that the CA1 spines susceptible to undergo mGluR-LTD are large-head mushroom spines that contain endoplasmic reticulum (ER) and often harbor a spine apparatus, whereas spines without ER are refractory to this plasticity (Holbro et al., 2009). Thus, we decided to assess the proportion of filopodia, stubby and mushroom shape spines but in principle we found no significant differences in the proportion of the different morphology categories among the experimental groups (Table 3). Nevertheless, a two-tail Z-score test for comparison of proportions from two populations showed p-values near significance when comparing proportions of filopodia in WT and WT RAPA groups (p-value = 0.0629) and between Ts1Cje and Ts1Cje RAPA groups (p-value = 0.0672), suggesting that prenatal treatment with rapamycin could increase the percentage of filopodia in both WT and Ts1Cje animals. Since, as stated above, a subpopulation of large-head, mushroom spines seem to be responsible for mGluR-LTD (Holbro et al., 2009), we measured the size distribution of the diameter for this particular type of spines. We found that the proportion of 0.5-0.7 μm mushroom spines was significantly reduced in Ts1Cje animals, compared to WT, and recovered in Ts1Cje RAPA mice, whereas the percentage of 0.7-0.9 μm mushroom spines was significantly increased in Ts1Cje mice and restored in Ts1Cje RAPA animals (Fig. 7). Thus, prenatal treatment with rapamycin normalizes the size distribution of mushroom type dendritic spines in Ts1Cje stratum radiatum neurons.

**Table 3.** Dendritic spine morphology comparison. The percentages of filopodia, stubby and mushroom spines present in secondary dendrites of apical stratum radiatum CA1 neurons in WT, Ts1Cje, and prenatally rapamycin-treated WT and Ts1Cje mice are indicated. Total number of spines (n) is also indicated for each experimental group.

| **Spine**<br>**Morphology** | **WT**<br>(n = 248) | **Ts1Cje**<br>(n = 151) | **WT**<br>**RAPA**<br>(n = 143) | **Ts1Cje**<br>**RAPA**<br>(n= 635) |
| --- | --- | --- | --- | --- |
| Filopodium | 14.9 | 13.2 | 22.4 | 19.7 |
| Stubby | 27.8 | 29.8 | 25.2 | 26.5 |
| Mushroom | 57.3 | 57.0 | 52.4 | 53.9 |

**Figure 7.**
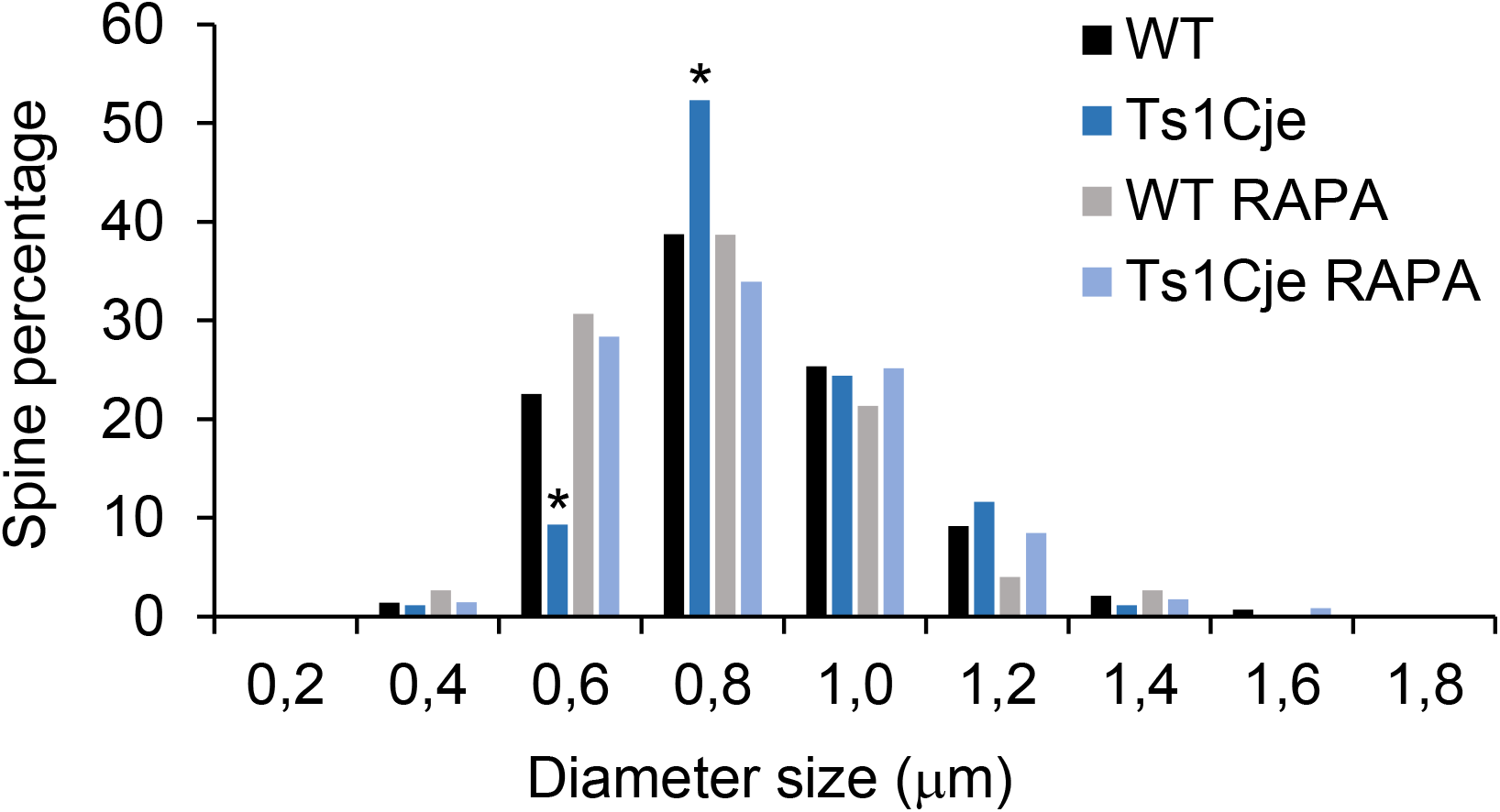
Frequency distribution of mushroom spines clustered by diameter size in CA1 stratum radiatum of untreated animals (WT and Ts1Cje) or mice treated with rapamycin prenatally (WT RAPA and Ts1Cje RAPA). Frequencies are shown as percentages. Center of the first and last bins in the histogram were automatically fixed (using GraphPad Prism software), and bin wide was set to 0.2 μm. The Z-score for two population proportions was calculated between WT vs. Ts1Cje, WT vs. WT RAPA and Ts1Cje vs. Ts1Cje RAPA for each histogram interval. Statistically significant p-values were obtained when comparing WT vs. Ts1Cje, and Ts1Cje vs. Ts1Cje RAPA, (as indicated with asterisks) in the following cases: 0.5 to 0.7 μm interval (bin center 0.6 μm): WT vs. Ts1Cje p-value = 0.011, Ts1Cje vs Ts1Cje RAPA p-value < 0.001; 0.7 to 0.9 μm interval (bin center 0.8 μm): WT vs. Ts1Cje p-value = 0.045, Ts1Cje vs Ts1Cje RAPA p-value = 0.002.

### Prenatal treatment with rapamycin normalizes mGluR-LTD in Ts1Cje hippocampus

In order to determine if the normalization of mushroom spine size distribution observed in Ts1Cje treated prenatally with rapamycin (Fig. 7) correlated with an effect on mGluR-LTD, we evaluated this plasticity in hippocampal slices of postnatal Ts1Cje mice treated prenatally with rapamycin (see Materials and Methods). Remarkably, as shown in Fig. 8, mGluR-LTD was normalized in Ts1Cje RAPA mice (91.55 ± 5.23% of baseline, n = 7 slices from 4 mice).

**Figure 8.**
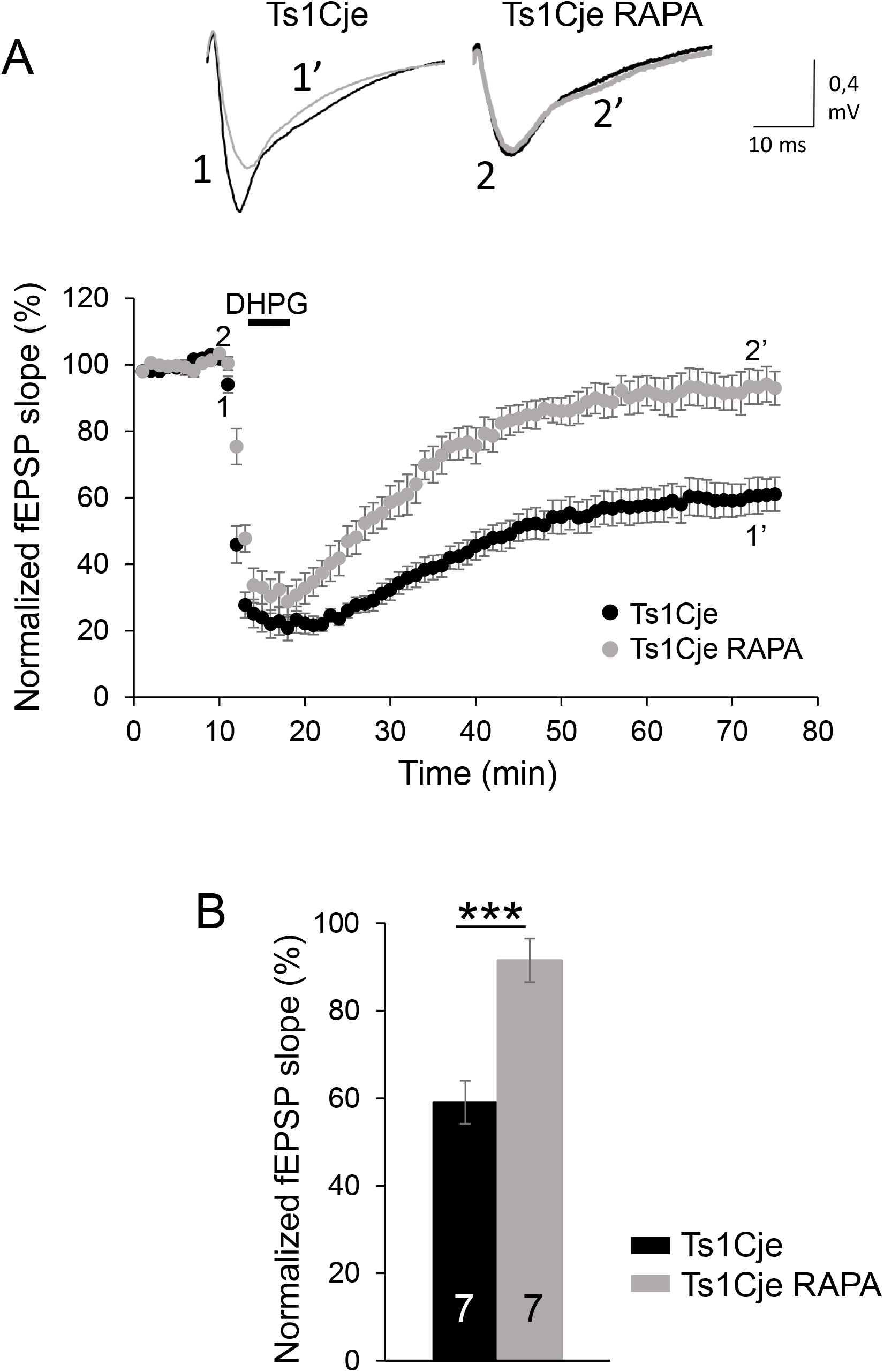
mGluR-LTD in hippocampal slices of Ts1Cje mice treated with rapamycin prenatally. (A) Time course of DHPG effects on field excitatory postsynaptic potentials (fEPSP) in Ts1Cje mice treated with rapamycin prenatally (Ts1Cje RAPA) and control Ts1Cje mice (same data as in Fig. 5). Upper insets: representative traces of a fEPSP before (1, 2) and after (1’, 2’) DHPG application in Ts1Cje (1, 1’; same data as in Fig. 5) and Ts1Cje RAPA (2, 2’) mice. (B) Quantification of the effects depicted in panel A. The error bars represent the SEM. The number of slices for each condition is indicated in the corresponding bar. P < 0.001 Student’s t-test.

## DISCUSSION

We have previously shown that the mTOR pathway is hyperactivated in Ts1Cje hippocampus (Troca-Marín et al., 2011). Afterwards, other groups reported mTOR hyperactivation in postmortem DS brain (Iyer et al., 2014; Perluigi et al., 2014). We have also found that rapamycin restored both BDNF-LTP and the persistence of LTM in the Barnes maze (Andrade-Talavera et al., 2015). We have here analyzed the performance of Ts1Cje animals in another hippocampal-dependent task, the NOR and found a clear memory index reduction in Ts1Cje mice, compared to WT. Furthermore, we found that rapamycin abolished NOR in WT mice, in agreement with previous reports (Jobim et al., 2012) but, remarkably, it improved NOR in Ts1Cje mice.

The characterization of the hippocampal synaptic proteome that we have here presented let to the identification of several affected pathways that could account for plasticity and memory deficits of Ts1Cje mice. Thus, two main functions were predicted to be altered in Ts1Cje hippocampus: mitochondrial function and synaptic plasticity (including LTD). Very interestingly, the synaptic levels of the proteins belonging to these pathways were normalized by rapamycin treatment of trisomic mice, strongly supporting the possible benefits of rapamycin treatment in the context of DS.

Mitochondrial dysfunction and increased oxidative stress have been previously found in the Ts1Cje brain (Shukkur et al., 2006). In fact, altered mitochondrial function has long been associated with DS (reviewed by Valenti et al., 2014; 2018; Izzo et al., 2018). It has been recently found damaged mitochondria linked to increased oxidative stress, reduced mitophagy and reduced autophagy, together with mTOR hyperactivation in fibroblasts from DS patients, (Bordi et al., 2019). The role of mTORC1 as regulator of both general autophagy and mitophagy induction after oxidative phosphorylation uncoupling is well established (Bartolome et al., 2017). Accordingly, pharmacological inhibition of mTOR using AZD8055, which inhibits both mTORC1 and mTORC2 (Chresta et al., 2010), restored autophagy and mitophagy in DS fibroblasts (Bordi et al., 2019). As mentioned before, we previously demonstrated hyperactivation of mTOR in the Ts1Cje hippocampus (Troca-Marín et al., 2011). Accordingly, we have here shown that autophagy is reduced, which could be related to the mitochondrial dysfunction that we detected in the proteomic analysis of Ts1Cje SNs.

Regarding synaptic plasticity, we found that mGluR-LTD is enhanced in Ts1Cje hippocampus. Many forms of synaptic plasticity have been characterized in the Ts1Cje and other DS mouse models (Siarey et al., 2005; Belichenko et al. 2007; Andrade-Talavera et al., 2015). mGluR-LTD has been previously studied in the Ts65Dn model but, in contrast to our results, a normal hippocampal mGluR-LTD, compared to WT, was found (Scott-McKean and Costa, 2011). Nevertheless, there are important differences between the Ts65Dn and Ts1Cje models that could explain this apparent contradiction. Thus, additional DS non-related trisomic genes exist in Ts65Dn (Duchon et al., 2011). The fact that 6-8 month-old mice were used in the Scott-McKean and Costa study while we used P21-30 mice could also explain the different results since the mechanisms of mGluR-LTD seem to be developmentally regulated (Moult et al., 2008).

mGluR-LTD is triggered by activation of group I mGluR (i.e., mGluR1/5) and relies on protein translation (Huber et al., 2000). Two main signaling cascades that regulate protein synthesis are engaged following mGluR1/5 stimulation: mTOR and ERK. Both pathways can stimulate cap-dependent translation at the initiation level. The relative contribution of these cascades to the protein synthesis necessary for mGluR-LTD is nevertheless unclear (for a review see Bhakar et al. 2012). Interestingly, it has been recently shown that mTORC2, but not mTORC1, is required for mGluR-LTD (Zhu et al., 2018). mTORC2 regulates actin cytoskeleton and, in fact, actin polymerization-depolymerization inhibition abolishes mGluR-LTD (Zhou et al., 2011). In any case, it seems clear that in wild-type conditions, rapid synthesis and degradation of FMRP is necessary for mGluR-LTD (Hou et al., 2006), which leads to transient local translation of key proteins involved in AMPAR internalization (i.e. GluA1 endocytosis), such as Arc/Arg3.1 (Park et al., 2008; Waung et al., 2008; Niere et al., 2012). Remarkably, and similarly to our results in Ts1Cje mice, it is well established that mGluR-LTD is exaggerated in FMRP knockout mice (Nosyreva and Huber, 2006), a model for Fragile X. However, Fragile X is due to loss of expression of FMRP and, in contrast, we found higher levels of FMPR in the Ts1Cje hippocampus, strongly suggesting a different cause for the enhanced hippocampal mGluR-LTD in trisomic mice.

Remarkably, it has been shown that the spines susceptible to undergo mGluR-LTD in the CA1 constitute a particular subpopulation of large-head mushroom spines with ER and spine apparatus, whereas spines without ER are refractory to mGluR-LTD (Holbro et al., 2009). We have here shown that the percentage of mushroom spines with a 0.7-0.9 μm size was notably increased in Ts1Cje CA1 dendrites, compared to WT (Fig. 7). Although we are not sure if this subpopulation corresponds to that described by Holbro and col., it is tempting to speculate that exaggerated mGluR-LTD in Ts1Cje hippocampus could be due to a higher proportion of ER-containing spines, susceptible to undergo mGluR-LTD. Excitingly, rapamycin treatment of pregnant dams normalized both the referred mushroom spine phenotype and mGluR-LTD in the Ts1Cje offspring.

mGluR stimulation leads to local inositol trisphosphate (IP3) receptor activation and calcium release from the ER (Holbro et al., 2009). Moreover, ryanodine receptors (RyRs) are particularly abundant in the spine apparatus (Sharp et al., 1993), an ER membrane specialization of stacked discs, which plays roles in local translation and calcium signaling (Jedlicka et al., 2008). mGluR-LTD induces trafficking from the ER to the synapse of GluA2, an AMPAR subunit that renders the receptor impermeable to calcium. This trafficking depends on IP3 and RyR mediated calcium release, and translation (Pick et al., 2017; Pick and Ziff, 2018). Interestingly, IPA of the Ts1Cje hippocampal SNs proteomic data predicted increased calcium signaling due to higher levels of RyR2, glutamate receptor subunits (including GluA2, GluA4, GluN1, GluN2B), plasma membrane calcium ATPases, and calcium voltage-gated channel subunits (Table 2). We think that the increased levels of these proteins could be a consequence of the higher percentage of the 0.7-0.9 μm mushroom spines in Ts1Cje hippocampus. Nevertheless, it should be noted that adult animals were used for proteomics while juvenile mice were used for electrophysiology experiments and spine morphology assessment. Thus, these correlations should be taken with precaution.

The molecular mechanism behind the mushroom spine morphologic phenotype in Ts1Cje hippocampal dendrites is unknown. Nevertheless, since it is recovered by rapamycin, mTOR signaling should be involved. We have previously found increased levels of both phospho-S6 (Ser235/236), a redout of mTORC1, and phospho-Akt (Ser473), a redout of mTORC2, in dendrites of Ts1Cje hippocampal neurons (Troca-Marín et al., 2011). Accordingly, either mTORC1 or mTORC2 (or both) could be involved. Interestingly, RICTOR, a canonical component of mTORC2, interacts with Tiam1, a Rac-1 specific guanine nucleotide exchange factor, to regulate actin polymerization (Huang et al., 2013). Tiam1 is encoded in the human chromosome 21 and it is in trisomy in Ts1Cje mice. Thus, Tiam1 overexpression in addition to mTORC2 hyperactivation could be relevant for the spine phenotype. Alternatively, the mushroom spine phenotype of Ts1Cje mice could be explained by mTORC1 hyperactivation and reduced autophagy since autophagy is required for developmental spine pruning. Moreover, it has been shown that in mouse models of autism and Fragile X that show mTOR hyperactivation, the activation of neuronal autophagy rescues the synaptic pathology (Yan et al., 2018; Tang et al., 2014).

In conclusion, we have here shown abnormal, exaggerated hippocampal mGluR-LTD in Ts1Cje mice, which correlates with an increased proportion of 0.7-0.9 μm mushroom spines in CA1 dendrites. Although the precise molecular/cellular mechanisms for these phenotypes remain to be dilucidated, prenatal treatment with rapamycin restored both morphologic and functional phenotypes, highlighting the therapeutic potential of rapamycin/rapalogs for correcting synaptic defaults in DS.

## ACKNOWLEDGMENTS

This work was funded by the Junta de Andalucía (Spain, Grant P12-CTS-1818), the Ministerio de Economía, Industria y Competitividad (Spain, Grant: SAF2015-65032-R), the Fondo Europeo de Desarrollo Regional (FEDER) and the Fondation Jérôme Lejeune (France). We thank Mariló Pastor (IBiS Proteomics Service; Instituto de Biomedicina de Sevilla, Spain) for technical help and advice, the Supercomputing and Bioinnovation Center (Universidad de Málaga, Spain) for providing us with access to the IPA tool, and Francisco J. Tejedor for critically reading the manuscript.

